# Bacterial DNA promotes Tau aggregation

**DOI:** 10.1101/786640

**Authors:** George Tetz, Michelle Pinho, Sandra Pritzkow, Nicolas Mendez, Claudio Soto, Victor Tetz

**Affiliations:** Human Microbiology Institute, New York, NY 10027, USA; Tetz Laboratories, New York, NY 10027, USA; Mitchell Center for Alzheimer’s disease and related brain disorders, Department of Neurology, University of Texas McGovern Medical School, Houston, TX 77030, USA

## Abstract

A hallmark feature of Alzheimer’s disease (AD) and other tauopathies is the misfolding, aggregation and cerebral accumulation of tau deposits. Compelling evidence indicates that misfolded tau aggregates are neurotoxic, producing synaptic loss and neuronal damage. Misfolded tau aggregates are able to spread the pathology from cell-to-cell by a prion like seeding mechanism. The factors implicated in the initiation and progression of tau misfolding and aggregation are largely unclear. In this study, we evaluated the effect of DNA extracted from diverse prokaryotic and eukaryotic cells in tau misfolding and aggregation. Our results show that DNA from various, unrelated gram-positive and gram-negative bacteria results in a more pronounced tau misfolding compared to eukaryotic DNA. Interestingly, a higher effect in promoting tau aggregation was observed for DNA extracted from certain bacterial species previously detected in the brain, CSF or oral cavity of patients with AD. Our findings indicate that microbial DNA may play a previously overlooked role in the propagation of tau protein misfolding and AD pathogenesis, providing a new conceptual framework that positions the compromised blood-brain and intestinal barriers as important sources of microbial DNA in the CNS, opening novel opportunities for therapeutic interventions.

## Introduction

The pathogenesis of certain neurodegenerative and autoimmune diseases is characterized by the misfolding and aggregation of proteins with prion-like properties, such as β-amyloid (Aβ) and tau in Alzheimer’s disease (AD), alpha-synuclein (α-syn) in Parkinson’s disease, TDP-43 and SOD1 in amyotrophic lateral sclerosis, and IAPP in type 2 diabetes [1-5]. Despite the fact that these proteins are associated with different diseases, they all accumulate in tissues in the form of toxic misfolded protein aggregates that have the capacity to spread among cells and tissues during the pathological progression [1-4]. Indeed, studies on tau, Aβ, and α-syn have shown that inoculation with tissue homogenates or misfolded proteins obtained from hosts afflicted with neurodegenerative diseases results in the induction of the disease pathology in the recipient cellular or animal models [1-4]. Moreover, in animals not genetically programmed to spontaneously develop the disease, pathological induction has reportedly resulted in completely de novo diseases that are more akin to infectious prions [6,7].

AD is a devastating degenerative disorder of the brain for which there is no effective treatment or accurate pre-clinical diagnosis. The major neuropathological changes in the brain of AD patients are neuronal death, synaptic alterations, brain inflammation and the presence of protein aggregates in the form of extracellular amyloid plaques, composed of Aβ, and intracellular neurofibrillary tangles (NFTs), composed of hyperphosphorylated tau [8]. For many years it was thought that Aβ aggregation was the most important pathological event in the disease, but recent findings suggest that tau hyperphosphorylation, misfolding and oligomerization might play a major role in AD pathogenesis [8,9]. Indeed, all AD clinical trials based on Aβ as a therapeutic target have failed so far [10]. In addition, the burden of neurofibrillary tangles correlates well with the severity of dementia in AD [8]. Finally, increasing evidence from *in vitro* and *in vivo* studies in experimental animal models provides a compelling case for tau as a promising therapeutic target [11]. The current view is that Aβ pathology is the primary driving force of the disease initiation, but this is accomplished by induction of changes in tau protein leading to the neurodegenerative cascade [8]. Tau aggregates do not accumulate only in AD, but are a common feature of several neurodegenerative disorders, termed tauopathies [12]. In other tauopathies, such as Progressive supranuclear palsy, Chronic traumatic encephalopathy, Corticobasal degeneration, Pick’s disease and Frontotemporal dementia, tau leads to neurodegeneration in the absence of amyloid plaques. Pathological tau is characterized by the formation of both intra- and extracellular misfolded forms, indicating that the seeding agent could be localized outside as well as within neurons [13,14].

The misfolding and aggregation of proteins into prion-like amyloid aggregates is not restricted to human diseases, but it has been observed in diverse organisms and in many cases leading not to disease, but actually to a beneficial activity for the cell [3,15]. Functional prion-like protein aggregates have been described in various organisms from bacteria to humans, and participate in a variety of functions including biofilm development, regulation of cytotoxicity, modulation of water surface tension, formation of spider webs, eggshell protection, enhancing HIV viral infection, modulation of melanin biosynthesis, regulation of memory formation, epigenetic factors and storage of peptide hormones [3,15]. It has been proposed that functional prion-like amyloid aggregates may promote disease-associated aggregates through a cross-seeding mechanism [3,16]. Interestingly, a recent study reported that bacterial amyloids may play a role in α-syn aggregation [17]. This is interesting considering that several lines of evidence indicate that intestinal bacteria play a key role in the pathogenic cascade of both PD and AD [18,19]. Supporting the hypothesis that microorganisms play role in AD pathogenesis, HHV-1, *Porphyromonas gingivalis, Candida spp., Escherichia coli, Chlamydophila pneumoniae*, and *Borrelia burgdorferi* have been detected in the cerebrospinal fluid (CSF) and postmortem brains of individuals with AD [20-25]. Moreover, cell wall components of C.*pneumoniae*, an obligate intracellular bacteria, and *E.coli*, which can act as a facultative intracellular parasite, have been found within neurons; pointing out that some of these bacteria could directly invade neurons [26,27].

The mechanisms and factors responsible for initiating the protein misfolding process in AD remain poorly understood. Various molecules have been shown to bind Aβ and tau and be able to initiate and/or promote protein misfolding and aggregation [28]. Among them, nucleic acids (including DNA and RNA) have been shown to bind with relatively high affinity to various misfolded proteins and in some cases to promote or stabilize the aggregates [29,30]. Extracellular bacterial DNA induces prion aggregation within microbial biofilm matrices and plays the critical role of converting bacterial amyloid into highly ordered, aggregated cross-β structures enabling unique structural properties of microbial biofilms [31,32]. A recent report identified the association between bacterial extracellular DNA and the formation of heat-resistant protein fractions within different proteins – even those lacking prion domains [33]. We hypothesize that DNA can act as an efficient promoter for protein misfolding in AD pathogenesis. To test this hypothesis, we focused on tau, an intracellular protein that has been shown to bind tightly to DNA and RNA and protect nucleic acids from degradation [34,35]. Moreover, in AD brains nucleic acids have been detected attached to NFTs and intracellular inclusions primarily composed of tau [36]. Using an in vitro tau aggregation assay, we observed that DNA derived from different sources induced a robust promotion of tau misfolding and aggregation. The effect was particularly strong by addition of bacterial DNA.

## Results

Tau misfolding and aggregation follows a seeding nucleation mechanism that can be modeled in vitro using purified recombinant tau protein incubated in the presence of heparin at physiological pH and temperature [37, 38]. For our experiments, we used full-length human tau protein containing 4 microtubule binding repeats (4R) and 2 N-terminal inserts (2N). The 2N4R tau is the largest form of tau and the most active in promoting microtubule assembly [12,13]. In most tauopathies the 4R tau is the predominant form of the protein in the aggregates [12,13]. Incubation of tau at a concentration of 50 µM in 10 mM HEPES, pH 7.4 containing 100 mM NaCl and 25 µM heparin at 37 °C with constant agitation led to the formation of amyloid aggregates, detectable by the fluorescence emission of thioflavin T (ThT) (Fig. 1A). ThT is an amyloid-binding molecule, which emits fluorescence when bound to the aggregates and is widely used to characterize the kinetic of amyloid formation [39]. Under these conditions, tau form ThT-positive aggregates (Fig. 1B, 1C, Supplementary Fig. S1) with a lag phase of around 15h (Fig. 1A). To measure the amount of aggregated tau, we centrifuged the samples and evaluated the tau signal in pellet and supernatant by western blot. As shown in figure 1C, most of the protein was recovered in the pellet and appears as a smear of high molecular weight bands. This result suggests that, under the conditions used the majority of tau was forming part of large aggregates. To use this aggregated material as seeds, we sonicated the preparation in order to generate seeding-competent short fibrils, as described in recent publications to produce tau preformed fibrils (tau-PFF) [40,41]. Analysis by transmission electron microscopy, revealed that this preparation contains short unbranched amyloid-like fibrils of different sizes and the expected width of ∼10 nm (Fig. 1D). One of the typical biological activities of tau aggregates is their ability to seed aggregation of monomeric tau. To test this property, we incubated tau at a lower concentration (20 µM), with less heparin (4 µM) and lower temperature (20 °C) in order to slow down spontaneous aggregation and observe a clear seeding activity (Fig. 1E). Addition of different quantities of PFF tau aggregates (sonicated fibrils) accelerated tau aggregation in a concentration-dependent manner (Fig. 1E). Indeed, the lag phase was directly proportional to the logarithmic amount of tau aggregates added as seeds (Fig. 1F). The lag phase was measured as the time in which aggregation begins, which is defined as the moment when the ThT fluorescence reaches a threshold of 40 fluorescence units.

**Figure 1.**
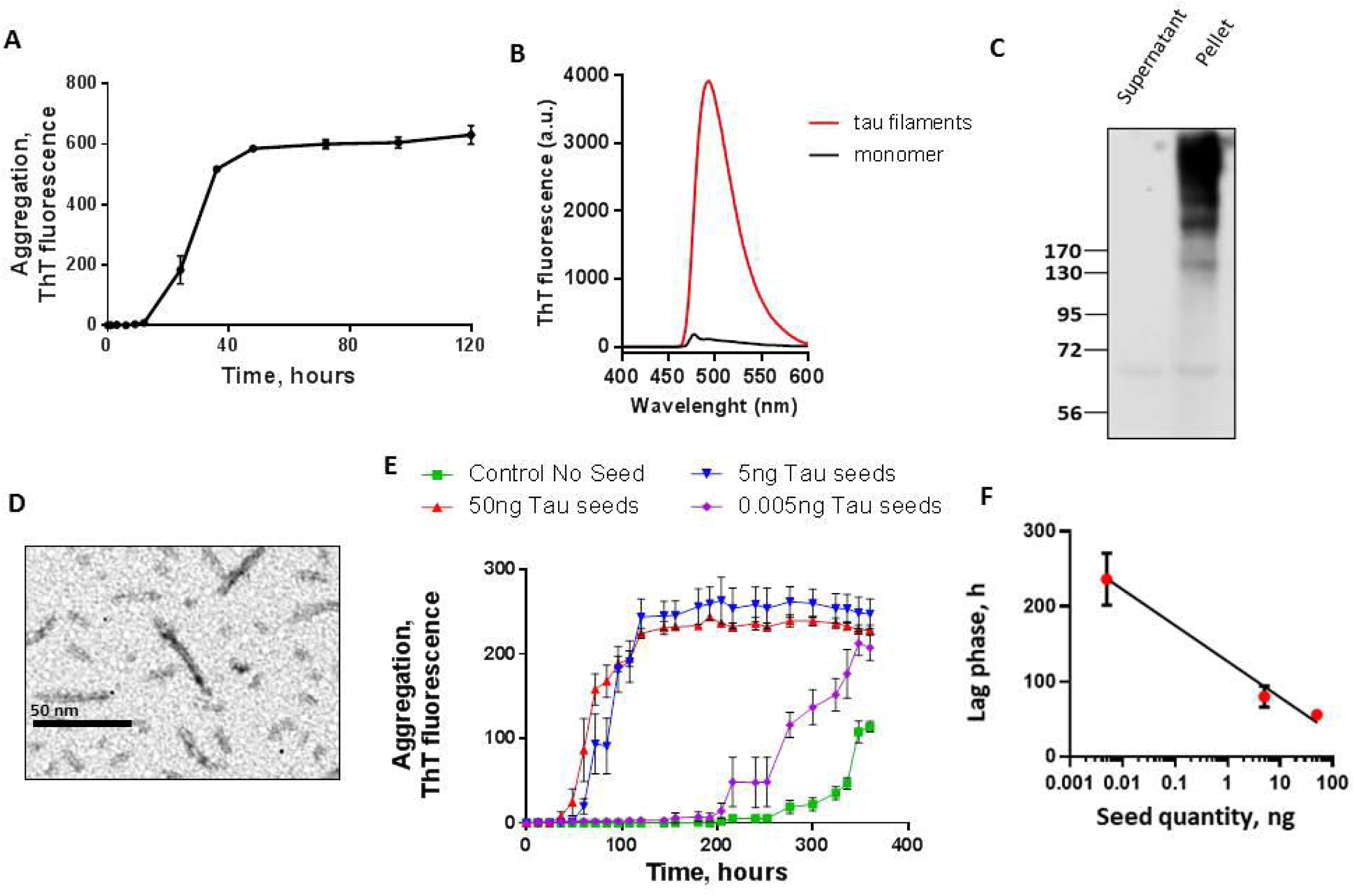
Tau seeding aggregation assay. **A**, Full-length Tau seeds were prepared by incubating tau monomer (50 µM) with 12.5 µM heparin in 10 mM HEPES pH 7.4, 100 mM NaCl for 5 days at 37°C with shaking. Aggregation was monitored by ThT fluorescence. **B**, Tau aggregates exhibit the typical ThT fluorescence spectrum with a maximum around 495nm when excited at 435nm. **C**, The aggregation state was further confirmed by sedimentation followed by western blot, showing that the majority of tau appeared in the pellet in the form of large molecular weight bands. **D**, the morphological characteristics of the tau aggregates were studied by transmission electron microscopy after negative staining with uranyl acetate. **E**, The Tau aggregation assay was performed on 96 well plates using 22 µM Tau monomer, 4.4 µM heparin, 10 µM Thioflavin T, using cyclic agitation (1 min shaking at 500 rpm followed by 29 min without shaking). Aggregation was followed over time by ThT fluorescence using a plate spectrofluorometer (excitation: 435; emmision: 485). Graph show the mean and SD of three replicates. **F**, Relationship between the quantity of tau oligomers and the Tau-PMCA signal (time to reach 50% aggregation).

Using this tau aggregation assay, we examined whether tau fibril formation could be promoted in the presence of extracellular DNA from various species including bacteria, yeast and human. For the experiments, monomeric tau was incubated with preparations containing 100 ng of DNA extracted from different bacterial species including *Pseudomonas aeruginosa* (PA), *Tetzosporium hominis* (TH), *Tetzerella alzheimeri* (TA), *Escherichia coli* ATCC 25922 (EC25), *Escherichia coli* ATCC 472217 (EC47), *Porphyromonas gingivalis* (PG), *Borrelia burgdorferi* (BB). We also incubated tau with the same amount of DNA extracted from *Candida albicans* (CA) and human samples. The results showed that DNA from various (but not all) bacterial species significantly promoted tau aggregation (Fig. 2A). Conversely, addition of eukaryotic DNA, such as from yeast or human cells, had a much lower effect in promoting tau aggregation. To compare the magnitude of the effect of these different DNA extracts on the kinetic of aggregation, we measured the lag phase, defined as the time at which aggregation begins, which is experimentally determined as the time when ThT fluorescence reaches a value >40 fluorescent units (equivalent to ∼2-folds the background levels after subtraction of the blank) [42]. Comparisons of the lag phases indicate that the largest promoting effect (shorter lag phase) was obtained in the presence of *Tetzerella alzheimeri, Escherichia coli* ATCC 25922, and *Escherichia coli* 472217 (Fig. 2B). Interestingly, *Tetzerella alzheimeri* (VT-16-1752 gen.nov, sp.nov) is a new species which was isolated from the oral cavity of a patient with AD [43]. Moderate promoting effect was observed with *Porphyromonas gingivalis* and *Borrelia burgdorferi*, and no significant effect was detectable for *Pseudomonas aeruginosa* and *Tetzosporium hominis* (Fig. 2B) [44].

**Figure 2.**
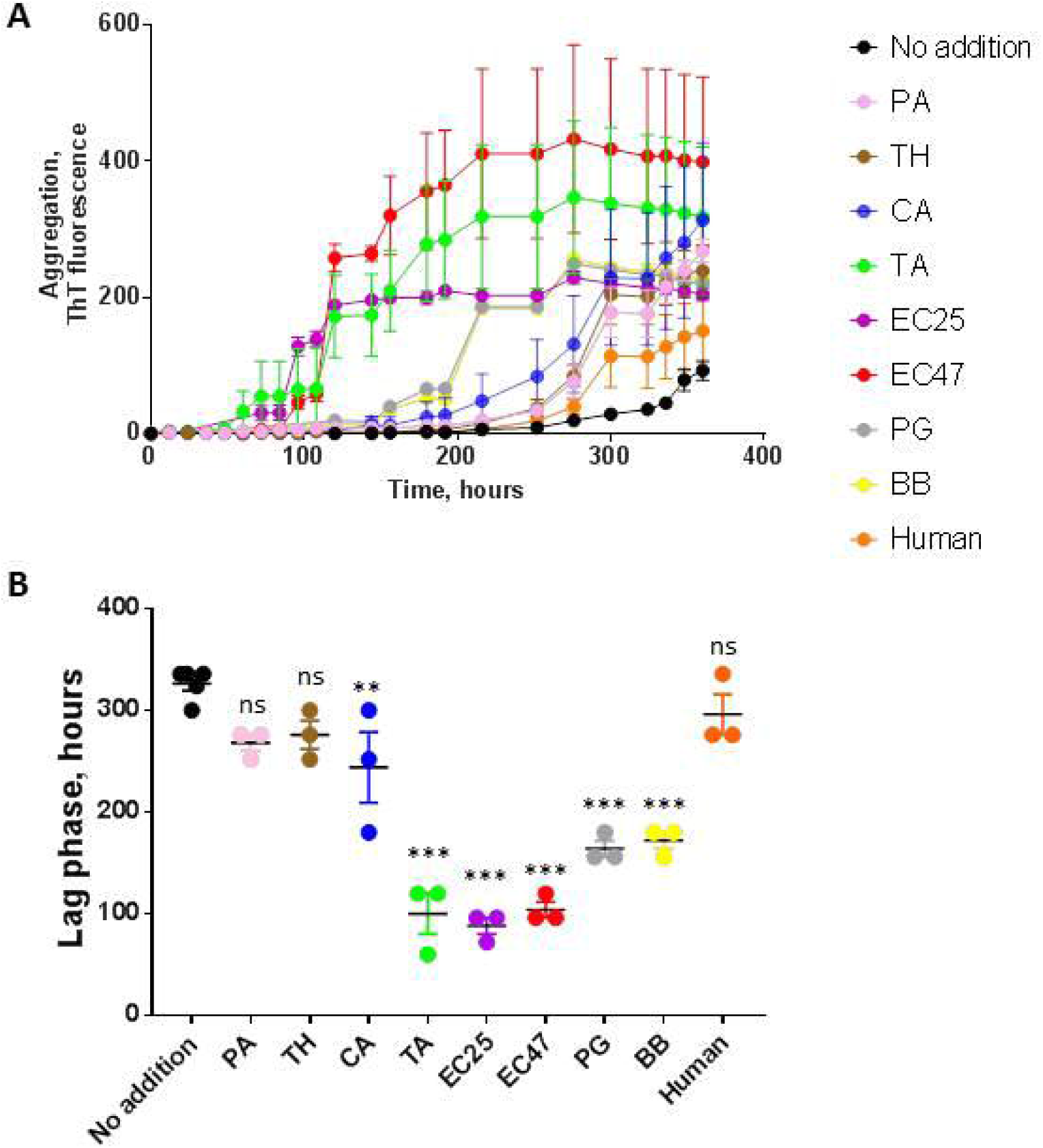
Effect of DNA extracted from diverse sources on tau aggregation. To study the effect of DNA on tau aggregation, monomeric tau (22 µM) under the conditions described in Fig. 1E, was incubated with preparations containing 100 ng of DNA extracted from different bacterial species including *Pseudomonas aeruginosa* (PA), *Tetzosporium hominis* (TH), *Tetzerella alzheimeri* (TA), *Escherichia coli* ATCC 25922 (EC25), *Escherichia coli* ATCC 472217 (EC47), *Porphyromonas gingivalis* (PG), *Borrelia burgdorferi* (BB). We also incubated tau with same amount of DNA extracted from *Candida albicans* (CA) and human samples. **A**, tau aggregation was monitored over time by ThT fluorescence. Data corresponds to the average ± standard error of experiments done in triplicate (except for control without seeds that was performed in quintuplicate). **B**, The lag phase, estimated as the time in which ThT fluorescence was higher than the threshold of 40 arbitrary units, was calculated for each experiment. The points represent the values obtained in each of the replicates. Data was analyzed by one-way ANOVA, followed by Tukey multiple comparison post-test. * P<0.05; ** P<0.01; *** P<0.001.

We then aimed to confirm whether the promoting activity of bacterial DNA is dose dependent. We found that DNA of *E.coli* ATCC 25922 and *P.gingivalis* at concentrations of 1000 to 10 ng significantly accelerated Tau aggregation relative to controls. The promoting activity of *E.coli* ATCC 25922 (Fig. 3) and especially *P.gingivalis* (Fig. 4) was lower than that of tau seeds. A dose dependent effect was more clearly observed only for addition of *P.gingivalis* DNA, perhaps because of the higher efficiency of *E.coli* ATCC 25922, which may require lower concentrations to observe a dose-dependency.

**Figure 3.**
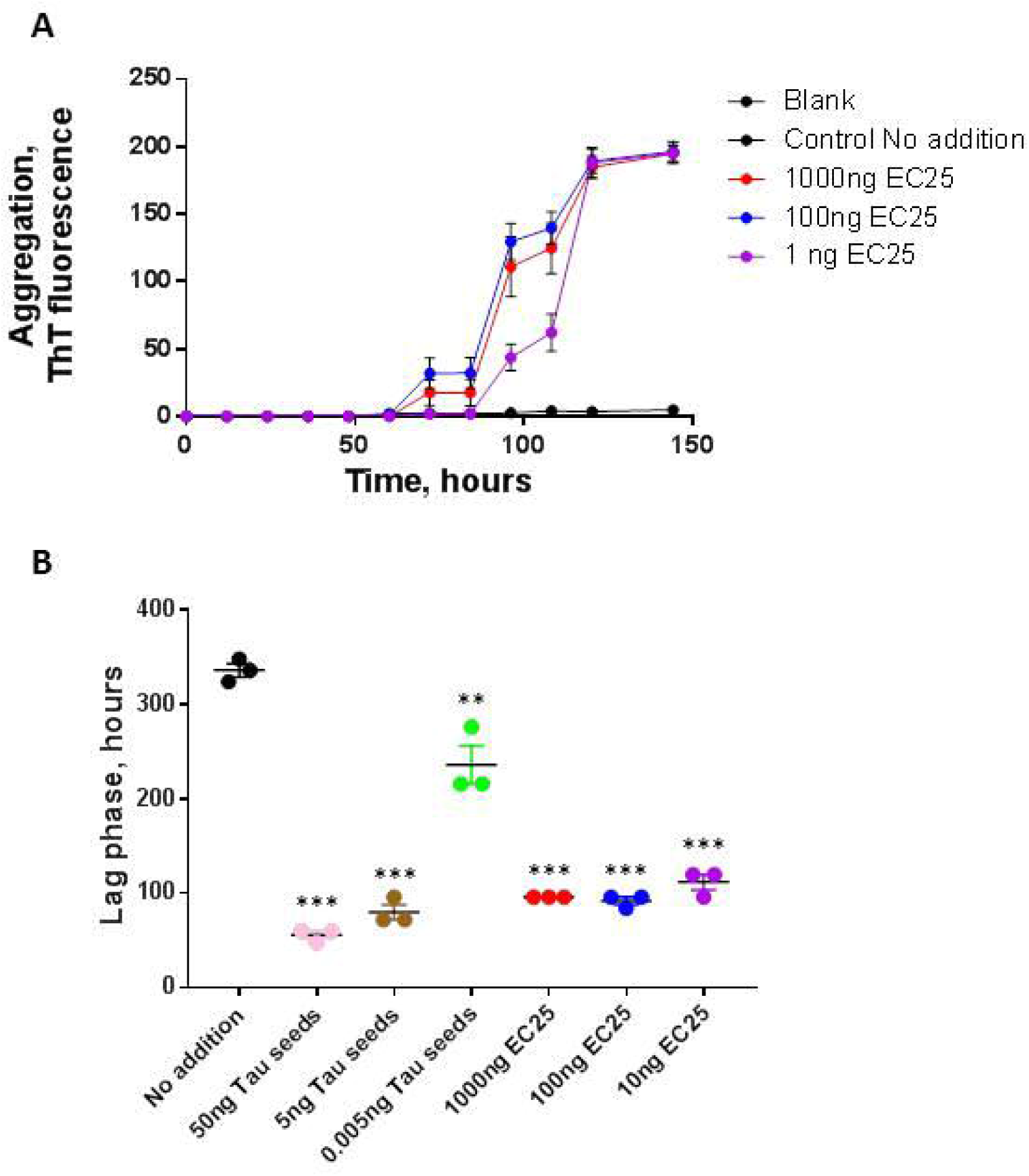
Influence of different concentration of *E. coli* ATCC 25922 DNA on tau aggregation. To study whether the promoting effect of *E.coli* DNA can be observed at different concentrations of DNA, we incubated monomeric tau under the conditions described above (Fig. 1E and Fig 2) with 1000, 100 and 10 ng of DNA extracted from *E. coli* ATCC 25922. **A**, tau aggregation was monitored overtime by ThT fluorescence. Data corresponds to the average ± standard error of experiments done in triplicate. **B**, The lag phase, estimated as the time in which ThT fluorescence was higher than the threshold of 40 arbitrary units, was calculated for each experiment. The points represent the values obtained in each of the replicates. Data was analyzed by one-way ANOVA, followed by Tukey multiple comparison post-test. * P<0.05; ** P<0.01; *** P<0.001.

**Figure 4.**
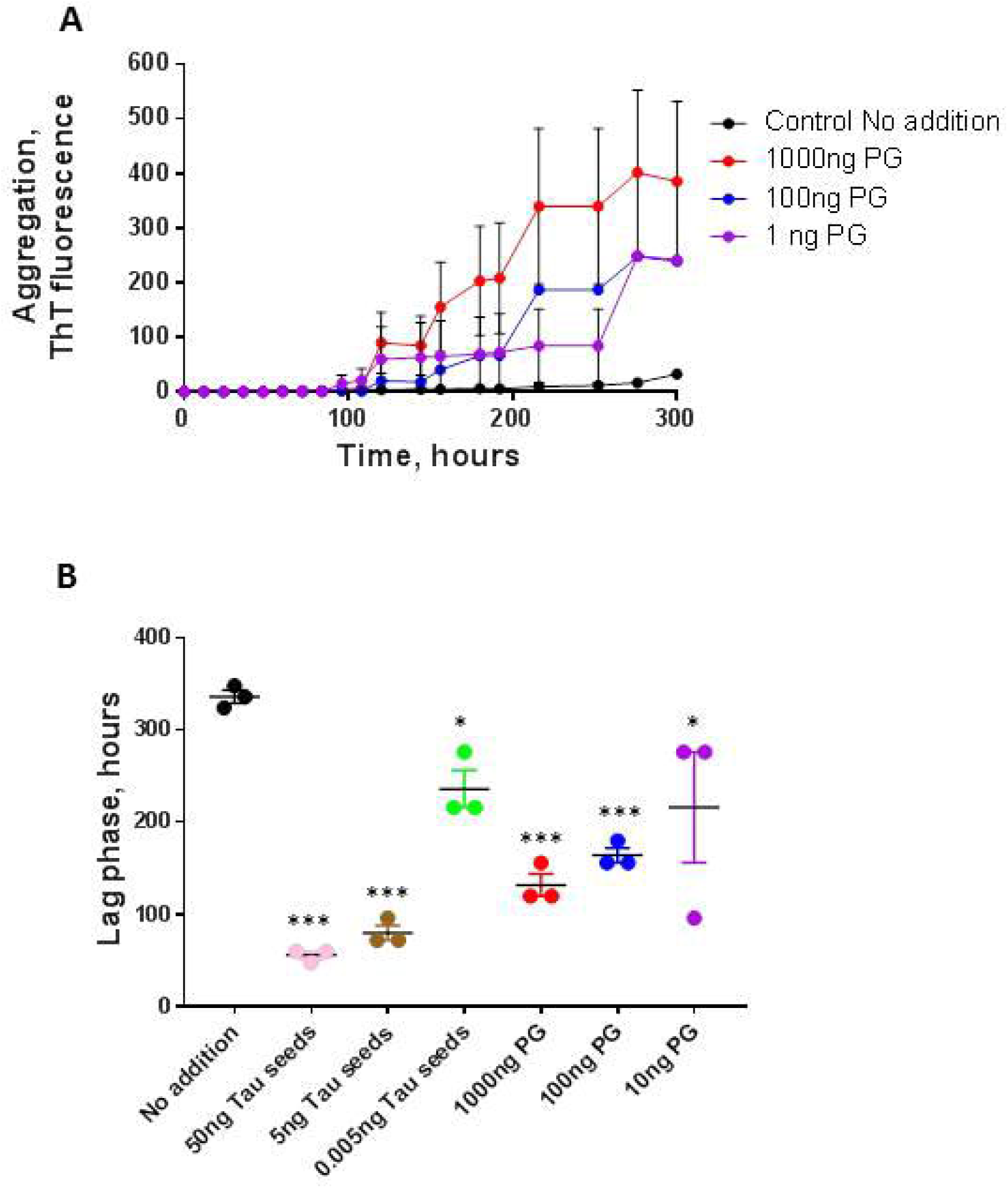
Dose-dependent effect of DNA from *Porphyromonas gingivalis* on tau aggregatio. Monomeric tau was incubated under the conditions described above (Fig. 1E and Fig 2) with 1000, 100 and 10 ng of DNA extracted from *P. gingivalis*. **A**, tau aggregation was monitored overtime by ThT fluorescence. Data corresponds to the average ± standard error of experiments done in triplicate. **B**, The lag phase, estimated as the time in which ThT fluorescence was higher than the threshold of 40 arbitrary units, was calculated for each experiment. The points represent the values obtained in each of the replicates. Data was analyzed by one-way ANOVA, followed by Tukey multiple comparison post-test. * P<0.05; ** P<0.01; *** P<0.001.

## Discussion

In addition to being one of the most devastating diseases of the 21st century, AD remains incurable. The cognitive symptoms and neurodegeneration appear to be mostly related to the extensive synaptic dysfunction and neuronal death observed in the brain [8]. In turn, neuronal lost and synaptic damage appears to be mediated by the progressive misfolding, aggregation and deposition of Aβ and tau proteins forming protein aggregates able to spread from cell-to-cell by a prion-like mechanism [1,3,4]. Genetics alone cannot account for the complex process of protein misfolding, aggregation and subsequent neurodegeneration observed in AD, particularly because the large majority of the cases are not associated to genetic mutations. Thus, it is likely that diverse environmental factors and age-related abnormalities play an important role on the initiation of the pathological abnormalities [45]. In this sense, various studies have shown that bacterial infection, as well as alterations in the intestinal microbiome may be implicated in the AD pathology [18, 46, 47].

Here we report the first evidence for the capacity of extracellular DNA from certain bacterial species to substantially promote tau misfolding and aggregation. The promoting effect of DNA on tau aggregation was observed in a wide range from 10 to 1000 ng. The use of these concentrations were informed by the range of cerebrospinal fluid DNA concentrations observed in patients with different diseases: 1 – 600 ng/mL [48-50]. The sources of bacterial and fungal DNA were selected based on the literature and personal data that showed associations of certain microorganisms with AD. Among the bacteria previously cultivated directly from the brains of patients with AD, or those whose components (such as nucleic acids, lipopolysaccharides, enzymes) were identified in the CSF, amyloid plaques, or brains of patients with AD, we used the DNA from *B. burgdorferi, P.gingivalis, C.albicans*, and *E.coli* [20, 20, 51-53].

Our data indicate that DNA from various, unrelated gram-positive and gram-negative bacteria significantly accelerated Tau aggregation. One of the best promoters was DNA from *E.coli* species, which is interesting for several reasons. First, Zhan et al demonstrated that some strains of *E. coli* (such as K99) were detectable immunocytochemically in brain parenchyma and vessels in AD patients more frequently compared to control brains [53]. Second, *E.coli* is known to share properties of facultative intracellular parasites and be localized within hippocampal neurons [54]; the latter finding is significant, as the hippocampus is the brain region greatly damaged in AD [55]. The intracellular localization of *E.coli* introduces unique possibilities regarding the interaction of bacterial DNA with tau proteins inside the neuron; e.g., the DNA can be secreted via transportation to the outer membrane or released following prophage induction and directly access the host neuron’s cytosol, where tau is normally present. Multiple lines of evidence indicate that the oral microbiome is implicated in AD development [23, 24, 46, 56, 57]. Orally localized bacteria may gain access to the brain via multiple pathways. As a peripheral chronic infection, periodontitis can trigger the onset of proinflammatory signaling cascades, weakening the blood-brain barrier (BBB) and resulting in direct CNS colonization by bacteria [46, 57, 58]. Moreover, the neurotropism of *spirochetal periopathogens* enables their spread through cranial nerves and propagation along the olfactory and trigeminal tracts [23, 24, 59]. The present study has found that the DNA of *P.gingivalis*, a causative agent of chronic periodontitis that has recently been isolated from the brain of people afflicted with AD [60] and is believed to be involved in the disease pathogenesis, triggers Tau misfolding. Moreover, DNA of other oral bacteria were also found to trigger Tau aggregation: *Tetzerella alzheimeri* VT-16-1752 gen. nov., sp. nov., belonging to *Brucellaceae* that was isolated from the prediodontal pocket of a patient with AD (complete genome sequence has been deposited in GenBank under the accession no. RQUW00000000) []. Notably, the DNA of *T. alzheimeri* [43] (together with that of *E. coli*) evinced the highest promoting activity relative to other bacteria.

The findings obtained in the present work indicate that DNA may play a previously overlooked role in the propagation of tau protein misfolding and AD pathogenesis, providing a new conceptual framework that positions the compromised blood brain and intestinal barriers as important sources of microbial DNA in the brain; indeed, altered gut permeability and disrupted BBB could precede AD development [61]. Moreover, we have recently described a clinical case showing improvement of cognition in an AD patient following therapy with deoxyribonuclease I enzyme, that cleaves cell-free DNA [62, 63].

In conclusion, the present study is – to the best of our knowledge – the first to outline the universal seeding effect of DNA isolated from different microorganisms on Tau protein aggregation. We found that DNA from various species of bacteria, some of which were previously identified in the CSF and brains of patients with AD, can lead to tau protein misfolding and aggregation, suggesting their potential role in the initiation and progression of pathological abnormalities responsible for AD.

Future studies should further investigate the possible role of DNA as an initial seeding factor for protein misfolding, including those associated with neurodegeneration, autoimmune diseases, and cancer. Moreover, subsequent studies should explore the targeting of DNA as a therapeutic strategy to prevent tau aggregation.

## Materials and Methods

### Sources and Procedures for DNA Extraction

Extracellular DNA was extracted from the matrix of *P. aeruginosa* ATCC 27853, *E. coli* ATCC 25922, *Escherichia coli* 472217, *Porphyromonas gingivalis, Borrelia burgdorferi Tetzerella alzheimeri* VT-16-1752, *Tetzosporium hominis* and *Candida albicans.*

All bacterial strains were subcultured from freezer stocks onto Columbia agar plates (Oxoid, UK) and incubated at 37 °C for 48 h, fungal strain was subcultured from freezer stocks onto Sabouraud dextrose agar (Oxoid, UK) and incubated at 30 °C for 48 h.

To extract the extracellular DNA, bacterial and fungal cells were separated from the matrix by centrifugation at 5000 g for 10 min at 4 °C. The supernatant was aspirated and filtered through a 0.2-μm-pore-size cellulose acetate filter (Millipore Corporation, USA). Extracellular DNA was extracted by using a DNeasy kit (Qiagen). Human genomic DNA (Roche Cat#11691112001) was purchased from Sigma (Sigma-Aldrich).

+ DNase I>>>>>>>>>>>>>>>>>>>

### Tau Expression and Purification

For these studies we used full-length Tau containing 4 microtubule-binding domains and 2 N-terminal inserts, with its 2 cysteines residues (C291, C322) replaced by serines [38] to prevent formation of covalent dimers and aggregates. The plasmid encoding Tau40 was kindly provided by Dr Martin Margittai. Expression and purification was done using previously described procedures [38].

Briefly, the plasmid was transformed into BL21 (DE3) Escherichia coli bacteria (New England BioLabs, catalog # C2527H) and grown overnight in 30 μg/ml kanamycin Terrific Broth (TB) at 37°C with agitation. The culture was then diluted 1:20 and grown until the optical density 600nm reached 0.6. One mM isopropropyl-β-D-thiogalactopyranoside (IPTG) was added to induce protein expression, and then the cultures were grown at 37°C with agitation for 6 hours. Bacteria were collected by centrifugation at 3,000xg and pellets stored frozen at −20°C until lysis.

Pellets were thawed and resuspended in 20 mM PIPES pH 6.5, 500 mM NaCl, with protease inhibitor cocktail complete (Roche) and sonicated with a 1/2” probe (S-4000, Misonix), then heated at 95°C for 20 minutes. Lysates were centrifuged at 15,000xg for 20 min at 4°C twice to remove cell debris. To precipitate proteins, ammonium sulfate (Sigma) was added at 55% w/v and incubated at room temperature for 1 hour with a magnetic stirrer. Precipitated protein was recovered by centrifugation at 15,000xg at room temperature, and pellets were stored at −20°C.

Tau protein was then purified by Cation Exchange Chromatography, pellets were dissolved in water (>18.2 MΩ cm) and then the solution was filtered through 0.2 µm filter. The sample was applied to a Hitrap SP HP column and eluted in a linear salt gradient (50-1000mM NaCl, 20 mM PIPES pH 6.5). The content of tau protein in the fractions was followed by SDS-PAGE and Blue Coomasie staining. Fractions containing tau protein were pooled and dialysed overnight 1:100 in 10 mM HEPES buffer pH 7.4, 100 mM NaCl.

Tau protein was concentrated in Amicon centrifugal filters 10 kDa MWCO and finally filtered through 100 kDa Amicon filter (Millipore) to remove pre-formed aggregates, aliquoted and stored at −80°C. Protein concentration was determined using BCA protein Assay (Thermo Scientific).

### Preparation of tau seeds

To prepare aggregated tau for seeding experiments, monomeric aggregate-free tau was incubated at a concentration of 50 µM, containing 25 µM heparin (Average MW 18,000, Sigma) in 10 mM HEPES buffer pH 7.4, 100 mM NaCl for 5 days at 37°C with constant shaking at 500 rpm in a thermomixer (Eppendorf). The formation of amyloid filaments was followed by Thioflavin T (ThT) fluorescence from samples taken from replicate tubes.

Tau filaments (2.3 mg/ml) were sonicated to prepare seeds (tau-PFF) by diluting the filaments to 0.1 mg/ml in 10 mM HEPES buffer pH 7.4, 100 mM NaCl and sonicated inside an Eppendorf tube in a floating rack using a microplate horn sonicator (S-400 Misonix) with settings of 30 seconds of total sonication time with pulses of 1 s ON −1 s OFF at Amp 30.

### Transmission electron microscopy (TEM)

An aliquot of 10 µl of reaction sample was placed onto Formvar-coated 200-mesh copper grids for 5 min, washed at least three times with distilled water, and then negatively stained with 2% uranyl acetate for 1 min. Grids were examined by electron microscopy (H-7600, Hitachi, Japan) operated at an accelerating voltage of 80 kV.

### Analysis of the amount of aggregated tau by sedimentation

To measure the quantity of tau remaining soluble and forming part of aggregates after incubation, we centrifuged samples at 100,000 xg for 1h at 4°C. The material in supernatant and pellet was analyzed by western blot after gel electrophoresis, using the anti-tau ab64193 antibody from Abcam.

### Tau aggregation assay

Solutions of 22 µM aggregate-free tau40 in 100 mM HEPES pH 7.4, 100 mM NaCl (200 µl total volume) supplemented with 4.4 µM heparin were placed in opaque 96-wells plates and incubated alone or in the presence of tau-PFF used as seeds to trigger tau aggregation or in the presence of distinct concentrations of DNA. Samples were incubated in the presence of 10 μM Thioflavin T (ThT) and subjected to cyclic agitation (1 min at 500 rpm followed by 29 min without shaking) using an Eppendorf thermomixer, at a constant temperature of 20 °C. At various time points, ThT fluorescence was measured in the plates at 485 nm after excitation at 435 nm using a plate spectrofluorometer.

### Statistical analysis

The significance of the differences in aggregation kinetics in the presence of different samples was analyzed by one-way ANOVA, followed by the Tukey’s multiple comparison post-test. To compare the effect of different samples, we estimated the lag phase, which corresponds to the time in which aggregation begins, defined as the moment when the ThT fluorescence reaches a threshold of 40 fluorescence units (around 2x the blank signal) after subtraction of the blank containing the buffer and ThT but not tau. The level of significance was set at P<0.05. Statistical tests were performed using the Graph Pad Prism 5.0 software.

## Figure Legends

**Supplementary Figure S1 . Full-length western blot**

Full-length western blot, showing that the majority of tau appeared in the pellet in the form of large molecular weight bands.

## Author Contributions

GT and VT designed experiments. MP and NM performed the experiments. SP and CS supervised the people performing the experiments. CS, VT and GT supervised data analysis, analyzed data and wrote the manuscript.

