## Supplementary figures and images for "Bacterial DNA promotes Tau aggregation"

### Supplementary figure 1

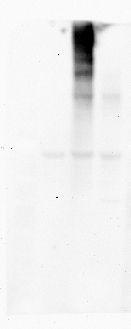
